# A novel *in vivo* system to study coral biomineralization in the starlet sea anemone (*Nematostella vectensis*)

**DOI:** 10.1101/2023.10.04.560932

**Authors:** Brent Foster, Fredrik Hugosson, Federica Scucchia, Camille Enjolras, Leslie Babonis, Will Hoaen, Mark Q. Martindale

**Affiliations:** The Whitney Laboratory for Marine Bioscience, Department of Biology, University of Florida, USA; Department of Ecology and Evolutionary Biology, Cornell University; School of Biosciences, Cardiff University, UK

**Keywords:** biomineralization, transgenesis, coral, *Nematostella vectensis*

## Abstract

Coral reefs are important for maintaining healthy marine ecosystems and are declining rapidly due to increasing environmental stresses. Coral conservation efforts require a mechanistic understanding of how these stresses may disrupt biomineralization, but progress in this area has been slow primarily because corals are not easily amenable to laboratory research. Some cellular characteristics of biomineralization are well characterized, such as the role of carbonic anhydrases, the polarized secretion of ions, and the secretion of “intrinsically disordered proteins” (IDPs) into extracellular microenvironments. We highlight how the starlet sea anemone (*Nematostella vectensis*) can serve as a tractable model to interrogate the cellular mechanisms of coral biomineralization. We have developed transgenic constructs using genes involved in biomineralization from several animal phyla that can be injected into *Nematostella* zygotes. These constructs are designed so translated proteins may be purified using TEV protease or Histidine tags to study their physicochemical properties. Using a fluorescent tag, we confirm ectopic expression of the coral biomineralizing protein SpCARP1 in live *Nematostella* embryos and adults and demonstrate via calcein staining that calcium ions co-localize with SpCARP1 in carbonate and calcium enriched seawater. Our findings suggest that SpCARP1 can induce the formation of amorphous calcium carbonate precursors in *N. vectensis*, consistent with its suspected role in the early stages of coral biomineralization. These results lay a fundamental groundwork for establishing *N. vectensis* as a novel *in vivo* system to explore the evolutionary and cellular mechanisms of biomineralization, improve coral conservation efforts, and even develop novel biomaterials.

**Graphical Abstract:** 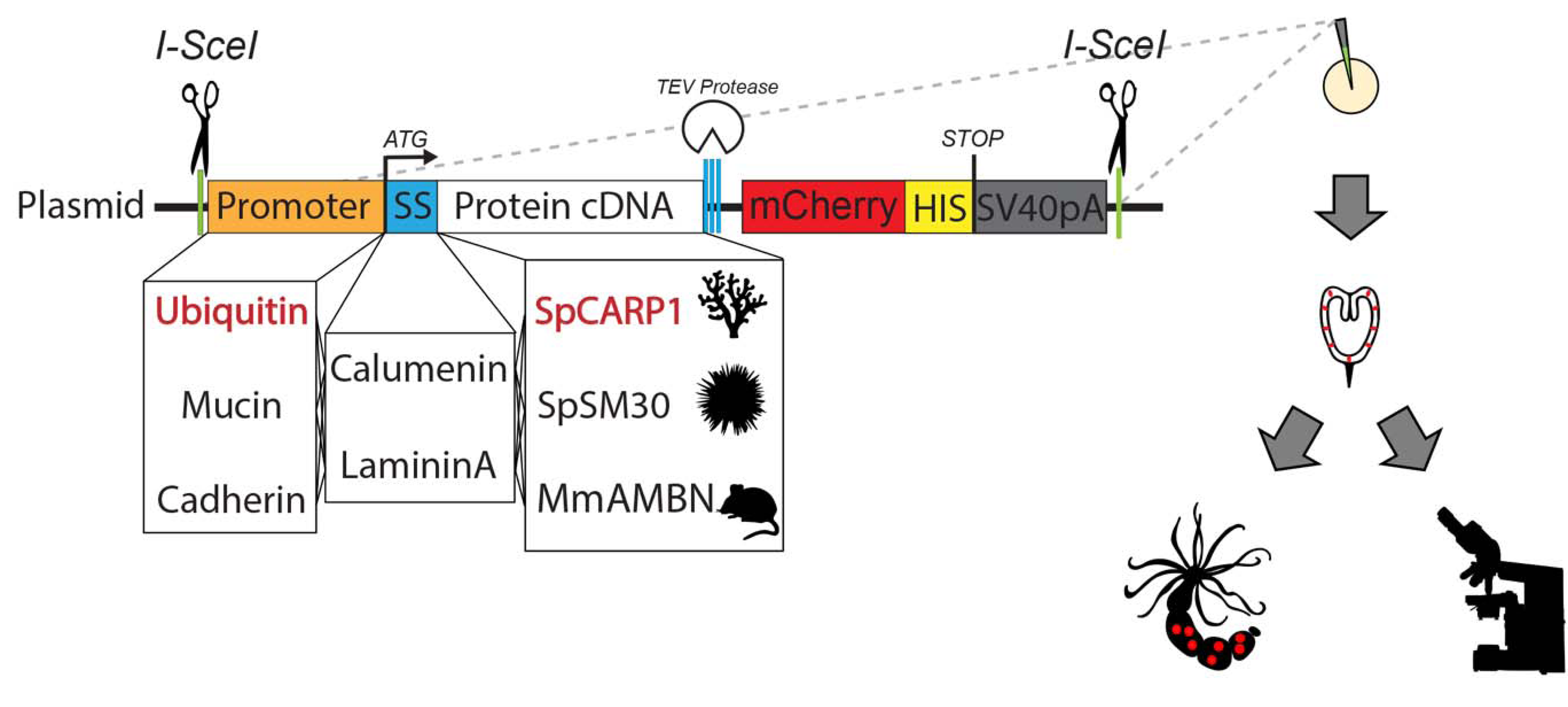

## 1 Introduction

Coral reefs represent some of the most biodiverse ecosystems on Earth^1–3^ and are necessary for maintaining healthy coastlines^4,5^. The backbone of these marine ecosystems are stony corals that, due to increased environmental stresses, are rapidly in decline^6^. Conservation efforts have been hampered, at least in part, by our limited understanding of the basic biology of corals and their ability to biomineralize and generate a diverse array of calcium carbonate skeletons that are susceptible to demineralization from changing ocean temperatures and acidification. Any meaningful effort to reverse the decline of corals requires a mechanistic understanding of 1) the molecular and biochemical processes of coral biomineralization and 2) how biomineralization is disrupted by environmental stresses. Unfortunately, efforts to probe the molecular basis of biomineralization in corals have proven difficult because of a general lack of genetic tools and difficulties culturing corals in laboratory settings.

Biomineralization is the production of inorganic minerals through biological mechanisms. This ability has evolved independently many times, resulting in unique structures such as bivalve shells^7–10^, sea urchin spicules^10–13^, and coral skeletons^14–16^. In marine organisms, the most studied mineralization pathways involve the absorption of Ca^2+ 17,18^ into cells expressing membrane-associated alpha carbonic anhydrases that convert CO_2_ to bicarbonate^19–22^. This results in the production of amorphous calcium carbonate (ACC) precursors that are stabilized to form crystal structures secreted into extracellular microenvironment^23–26^.

Energetically favorable conditions for biomineralization can arise spontaneously and rapidly, suggestive of a mechanism by which ACC biomineralization could have evolved independently through the use of non-homologous proteins with similar physicochemical characteristics that result in similar mineralized materials^23,26,27^. Secreted proteins within mineralizing cells have been shown to catalyze nucleation^28,29^ and/or interact with ACC precursors to provide stability as mineralizing tissue becomes more structured and complex^13,26,30–32^. These proteins are considered to be “intrinsically disordered” (IDP) because they have no set tertiary structures^33–36^. Although biomineralizing species may not share homologous IDPs, many of these proteins contain similar properties such as highly acidic residues^8,37–40^ and post-translational modifications that modify their folding and biomineralizing activity^32,40,41^.

Existing biomineralization models have distinct advantages and disadvantages. Bacterial expression systems help clarify the role of carbonic anhydrases in biomineralization^42–44^, yet they are unable to modify proteins endogenously after translation and therefore cannot be used for elucidating the role of post-translational modifications in biomineralization. Marine invertebrates offer several advantages to *in vivo* assays of biomineralization. Sea urchins are useful as developmental models to understand the dynamics of spicule growth during embryonic skeleton formation^32,45^ and syncytial mineralization ^46^. Mollusks can be used to test the effects of novel, taxon-specific proteins on shell formation^35,38^. Corals are useful for characterizing how matrix proteins stabilize biominerals in extracellular microenvironments^14,26^. In each of these *in vivo* systems, mechanistic studies of the dynamic processes of biomineralization can be difficult to interpret due to the complexity of interacting biomineralizing processes^47^.

Here, we present the starlet sea anemone (*Nematostella vectensis*) as a model for studying the dynamic processes of biomineralization. Despite being in the same class (Anthozoa) as scleractinian corals, *N. vectensis* does not naturally mineralize, eliminating potential confounding factors of interacting mineralization reactions^47^. Comparative genomics reveals that *N. vectensis* retains much of the molecular machinery believed to be necessary for biomineralization, including carbonic anhydrases^48^. *N. vectensis* is a powerful developmental model that can easily produce thousands of embryos on demand with simple light and temperature cues. Embryos are easy to microinject, and many techniques for manipulating gene expression are already well established in *N. vectensis*^49^, making this organism well-suited for investigating gene function during biomineralization.

In this article, we show that *N. vectensis* can express transgenic proteins involved in biomineralization in other taxa and present a novel *in vivo* system to evaluate the ability of IDPs to sequester calcium ions, enabling future studies to assess the role of these proteins in biomineral deposition.

## 2 Materials and Methods

### 2.1 Animal Culture

Adult *Nematostella vectensis* were maintained in 1/3X filtered seawater (FSW) diluted in deionized water and spawned following protocols as described previously^50–52^.

### 2.2 Molecular cloning and In vitro mRNA Transcription

As a proof-of-principle, we focused on a Coral Acid-Rich Protein (CARP) from a stony coral (*Stylophora pistillata*). SpCARP1 is a membrane-associated IDP that binds Ca^2+^ ions to induce CaCO_3_ precipitation and is believed to initiate biomineralization in *S. pistillata*^29^. To highlight the potential and versatility of our system for studying other forms of biomineralization, we also developed transgenic constructs for expressing proteins involved in the formation of sea urchin spicules (*Strongylocentrotus purpuratus*: SpSM30) and mice teeth (*Mus musculus:* Ameloblastin, MmAMBN).

SpCARP1 (KC148537) cDNA was synthesized by IDTDNA Inc. (idtdna.com). SpSM30 (NP_999766.1) cDNA was first codon optimized using Codon Optimization OnLine (COOL)^53^ and synthesized by IDTDNA Inc. (idtdna.com). MmAMBN cDNA (NM_001303431.1) was ordered from Genscript (New Jersey, USA; clone ID: OMu67099). Primers for SpSM30 and MmAMBN (**Supplementary Table 1**) were designed with Primer3^54^. All cDNA was cloned in frame into the pCS2+8CmCherry vector (Addgene Plasmid #34935) using AscI (NEB #R0558) and ClaI (NEB #R0197) cut sites. The SpCARP1 insert was synthesized as a gene fragment by Twistbioscience (Twistbioscience.com) and consisted of flanking restriction sites, a Kozak sequence optimized for invertebrates (AAAAAA)^55^, putative signal sequences native to *N. vectensis* (Calumenin: v1g117044 or Laminin A: v1g248148) replacing the predicted signal sequence in the SpCARP1 cDNA. A linker sequence (GGATCCGCTGGCTCCGCTGCTGGTTCTGGCGAATTC)^56^ and TEV protease recognition site were included in SpCARP1 and SpSM30 inserts. *Nematostella* signal sequences were predicted using SignalP^57^. mRNA was *in vitro* transcribed from linearized plasmids following the protocols for the Invitrogen mMessage mMachine SP6 Transcription Kit (Invitrogen AM1340) and purified using the MEGAclear Transcription Clean-Up Kit (AM1908). See also **Supplementary Table 1**.

### 2.3 Isolation of Promoter DNA Sequences

In order to express engineered proteins at distinct times and in specific cell types, we cloned putative promoter sequences upstream of the transcriptional start sites for *Nematostella Ubiquitin* (v1g217964) and *Mucin* (v1g203270) genes (see **Supplementary Table 2** for coordinates and primers used for cloning promoter sequences). Sequences were identified using the *Nematostella vectensis* genome 1.0^58^ and amplified from gDNA extracted from whole embryos or adult tentacle clips using standard PCR procedures. To initially test for promoter activity, DNA fragments were cloned into the pNvT-MHC::mCherry vector (Addgene #67943) using PacI (NEB #R0547) and AscI (NEB #R0558) sites, thereby replacing the myosin heavy chain (MHC) promoter. Confirmed plasmids were prepared following the protocol for the GeneJET Miniprep kit (ThermoFisher cat. #K0503). Sequences were confirmed via standard Sanger sequencing (Psomagen.com). When later cloned into pCS2+8CmCherry vector (see next section), the promoter sequences were cut out using SpeI (NEB #R3133) and AscI (NEB #R0558) sites (see **Supplementary Figure 1**).

### 2.4 Generation of expression constructs for Transgenesis

The software programs Serial Cloner V2.6 and Geneious Prime 2021.2.2 (https://www.geneious.com) were used to design transgenic constructs. The inserts were first cloned in frame into a pCS2+8CmCherry vector (Addgene Plasmid #34935) using AscI (NEB #R0558) and ClaI (NEB #R0197) sites. Promoter sequences were then inserted upstream using SpeI (NEB #R3133) and AscI (NEB #R0558) sites. Finally, fragments containing promoter and fusion protein segments were digested and cloned into the pKHR4 vector (Addgene #74592) using SpeI (NEB #R3133) and NotI (NEB #R0189) sites. The pKHR4 vector contains I-SceI endonuclease recognition sites flanking the multiple cloning site that was replaced with our inserts.

### 2.5 Microinjection

Fertilized eggs were prepared for microinjection as described previously^50^. Plasmids were incubated with 10X Cutsmart buffer and yeast I-SceI endonuclease (NEB #R0694) at 37ºC for approximately 30 minutes prior to injection and then mixed with either Rodamine Green or Alexa488 conjugated Dextran (0.2 mg/ml final concentration). Plasmids were injected in a final concentration of approximately 25 ng/μl. mRNA was diluted in nuclease-free water and mixed with nuclease-free Rodamine Green Dextran (0.2 mg/ml) and injected in final concentrations between 100 – 300 ng/μl.

### 2.6 Fixation and Confocal Microscopy

Animals were either live-imaged or fixed 24 hours post fertilization (hpf), 96 hpf, or 1-week postinjection as previously described^59,60^. Live animals were mounted in 1/3X FSW. Fixed animals were then washed in PBS-Tween, stained for DAPI and Alexa488 Phalloidin, and mounted on glass slides in either 80% glycerol or PBS. All animals were imaged on a Zeiss Imager. Z2 or a Zeiss 710 laser scanning confocal microscope. Confocal images were Z-stacked with max intensity in FIJI^61^ to show fluorescent signal.

### 2.7 Water Enrichment and Calcein Incubation

For both the non-enriched and enriched 1/3X FSW, temperature and pH (NBS scale) were measured using a pH/ATC electrode (Thermo Fisher Scientific, Waltham, USA), calibrated using pH 4, pH 7, and pH 10 buffer solutions (Thermo Fisher Scientific, Waltham, USA). Salinity was measured using a digital refractometer (Milwaukee Instruments, Rocky Mount, USA). Measurements of total alkalinity (TA) were performed using an alkalinity test kit based on drop count titration (sulfuric acid) (Hach, Loveland, USA). Parameters of seawater carbonate system were calculated from pH, TA, temperature, and salinity using the CO2SYS package^62^ with constants from^63^ as refit by^64^ (see **Supplemental Table 3)**.

The concentration of calcium and carbonate ions regulate the thermodynamic driving force that determines the precipitation of calcium carbonate in biomineralizing animals^47,65^. To replicate biomineralization-favorable conditions, we incubated 1-month-old *N. vectensis* injected with transgenic SpCARP1 constructs in either 10mM CaCl_2_, 10 mM NaHCO_3_, or 10mM CaCl_2_ + 10mM NaHCO_3_ in 1/3X FSW for 1 hour in a cell culture petri dish (5 mL). Polyps were then transferred to a new dish and incubated for another hour in a Calcein Blue solution (2.6 μM; Sigma 54375-42-2). Polyps were rinsed for 30 minutes in 1/3X FSW, then immobilized by adding 7.14% MgCl2 before imaging with a Zeiss 710 confocal microscope. Samples were observed with the mCherry red fluorescent filter (range 415–735nm) and the DAPI blue fluorescence filter (range 410–495nm) using 40X magnification. All imaging settings were kept constant between the samples. Images were acquired with the ZEN 2011 software (v14.0.0.0; Zeiss, United States) and processed in FIJI^61^.

### 2.8 Single Cell Dissociations

Injected embryos were dissociated 24 hours post injection in 1/3X Ca2+/Mg2+-free and EDTA-free artificial seawater as previously described^66^. Dissociated cells were incubated for 1 hour in 1:5000 CellMask (Fisher Scientific C37608), then washed two times in the dissociation media. Cells were water-immersed and imaged on a Zeiss Imager.Z2 at 40X magnification.

## 3 Results

### 3.1 Plasmid constructs are adaptable for targeted and stable transgenesis

*N. vectensis* embryos grow into swimming planulae within 48 hours post fertilization (hpf), settle, then develop into small polyps in about a week when kept at room temperature (25ºC) (**Figure 1A**). We also injected zygotes with a putative ubiquitin promoter driving mCherry fluorescent signal and show broad expression in planulae (**Figure 1B–B’**) and small polyps (**Figure 1C–C’**). We also designed plasmid vectors to incorporate other putative promoters endogenous to *N. vectensis*, as well as native signal sequences, driving expression of IDPs involved in biomineralization (**Figure 1D**). We could not detect any visible difference with plasmid constructs containing signal sequences (SS) native to *N. vectensis* or those present in non-native cloned constructs. As such, the remainder of our data makes no distinction of whether constructs contain SS endogenous to *N. vectensis* or cloned sequences.

**Figure 1.**
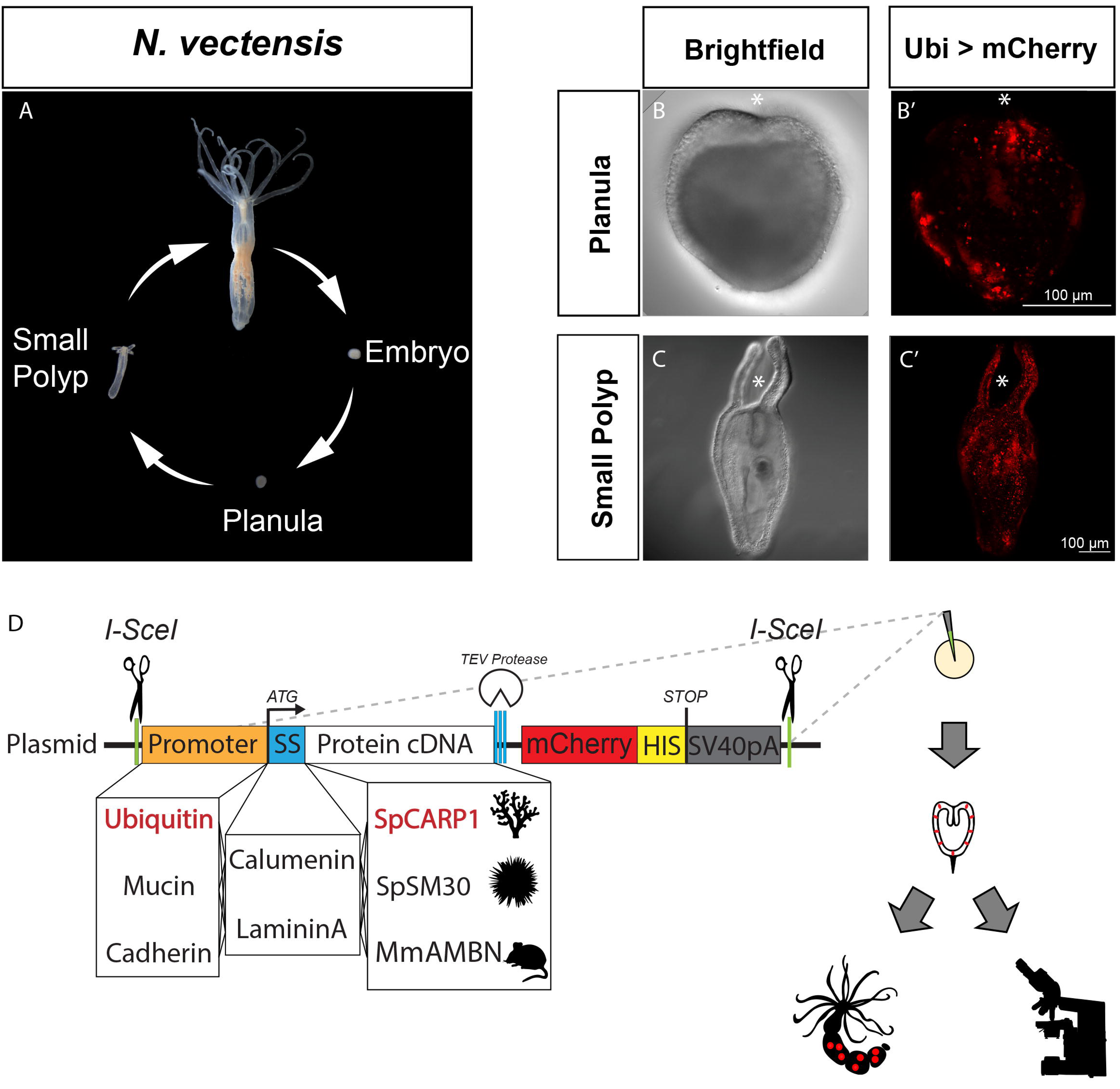
Putative promoters are sufficient to drive stable expression of mCherry. Life cycle of *N. vectensis* (A). Brightfield and Max projections showing the ubiquitin promoter driving expression of mCherry in live planula (B– B’) and small polyps (C–C’). Plasmid construct containing native promoters, signal sequences (SS), and proteins involved in biomineralization of coral (SpCARP1), sea urchin spicule (SpSM30), and mouse tooth enamel (MmAMBN), as well as the general workflow including microinjections, rearing of animals with incorporated transgene and evaluation of fluorescent mCherry signal with confocal microscopy (D). Asterisk = oral pole

Animals injected with the constructs containing the ubiquitin promoter driving expression of SpCARP1 exhibit transient expression as early as 24 hpf. By the planula stage, mCherry signal is broadly detected in both endoderm and ectoderm (**Figure 2A**). Expression expands into the body column and tentacles of developing small polyps (**Figure 2B**), with the strongest signal in scattered ectodermal cells (see arrows in **Figure 2A** and **2B**). SpCARP1::mCherry signal persists when cells are dissociated (**Supplementary Figure 3A–B**). Within 24 hours of dissociation, cells form aggregate clumps and maintain fluorescent signal (**Supplementary Figure 3C–D**). The mucin promoter drives expression of SpCARP1 within 48 hpf in aboral ectoderm of developing planulae in characteristic scattered secretory gland cells (**Figure 2C**). Strong mosaic signal expands into the body column and tentacles of small polyps in what appears to be glandular cells (**Figure 2D**; see arrows).

**Figure 2.**
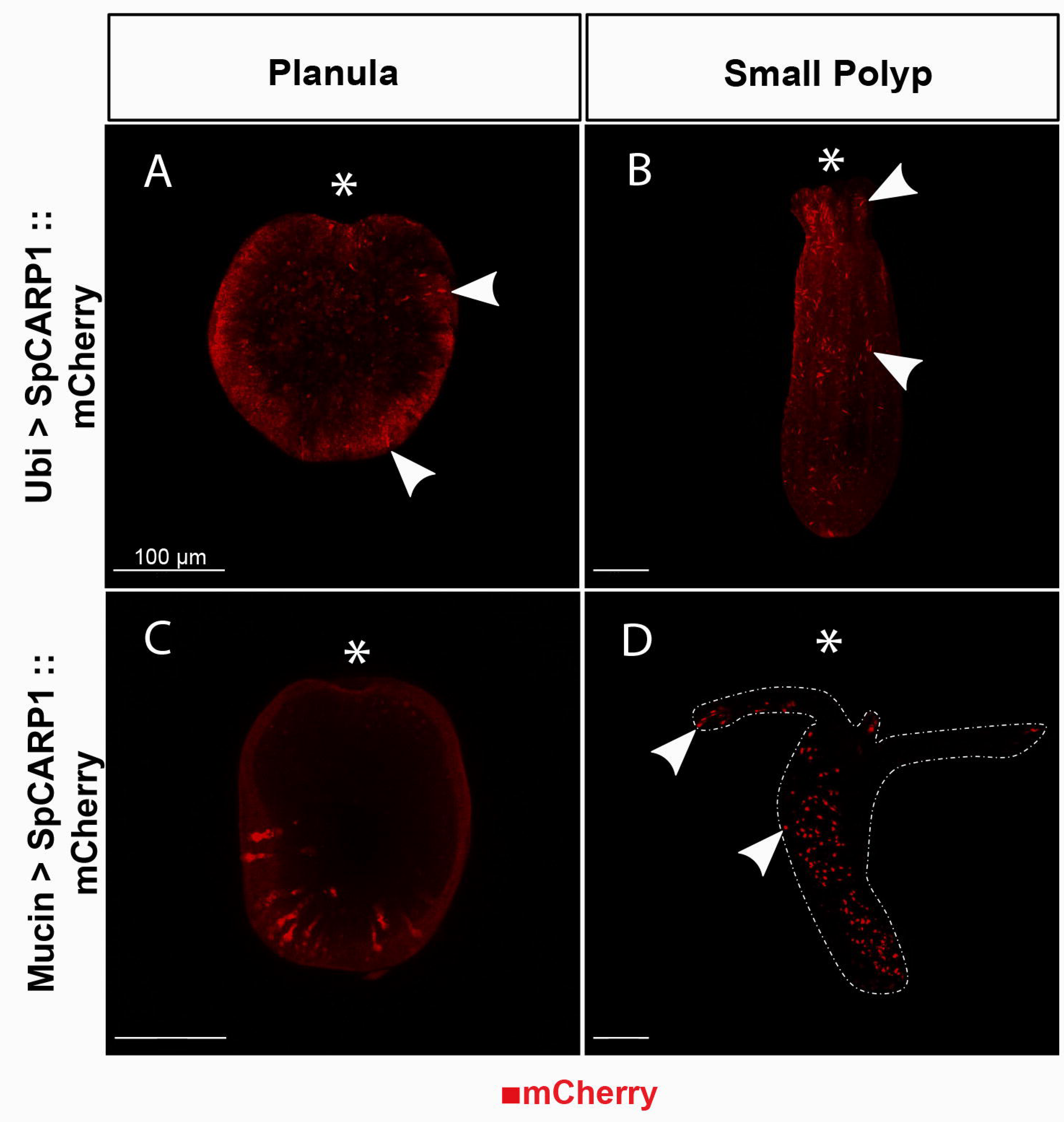
CARP1::mCherry expression can be driven by endogenous Nv promoters. Ubiquitin promoter drives broad expression in ectoderm in live planulae (A) and small polyps (B), with the strongest signal in scattered ectodermal cells (see white arrowheads). Mucin promoter drives expression in secretory cells in fixed planula aboral ectoderm (C) and throughout the body column and tentacles of small polyps (D). Asterisk = oral pole. All scale bars = 100μm.

### 3.2 SpCARP1 preferentially co-localizes with calcein in the tentacles of polyps

We imaged live transgenic polyps to observe the pattern of expression of SpCARP1 and the potential colocalization of the protein with calcium ions, suggestive of biomineralization-related activity. Limited calcein signal is also present in WT controls (**Figure 3A–F’**), indicating that the fluorescent dye binds to calcium ions naturally present in the organism. In transgenic polyps, noticeable mCherry fluorescence is localized primarily at the tip of the tentacles and sparse regions along the tentacle cavity (**Figure 3G–I’**; see white arrows). The mCherry signal is also present around the oral pole. Co-localization of calcein and mCherry fluorescence in the tentacles is evident in the endoderm of the tentacle cavity (**Figure 3G–I’**) and in sparse regions surrounding the mouth (**Figure H**). Along the body and in the aboral end, the mCherry fluorescence is prevalent in the endoderm and the gastrovascular cavity, whereas the calcein signal is mostly localized to the ectoderm (**Figure 3J–L’**). This pattern suggests that the calcium-binding activity of SpCARP1 is mostly concentrated in the tentacles of *N. vectensis* polyps.

**Figure 3.**
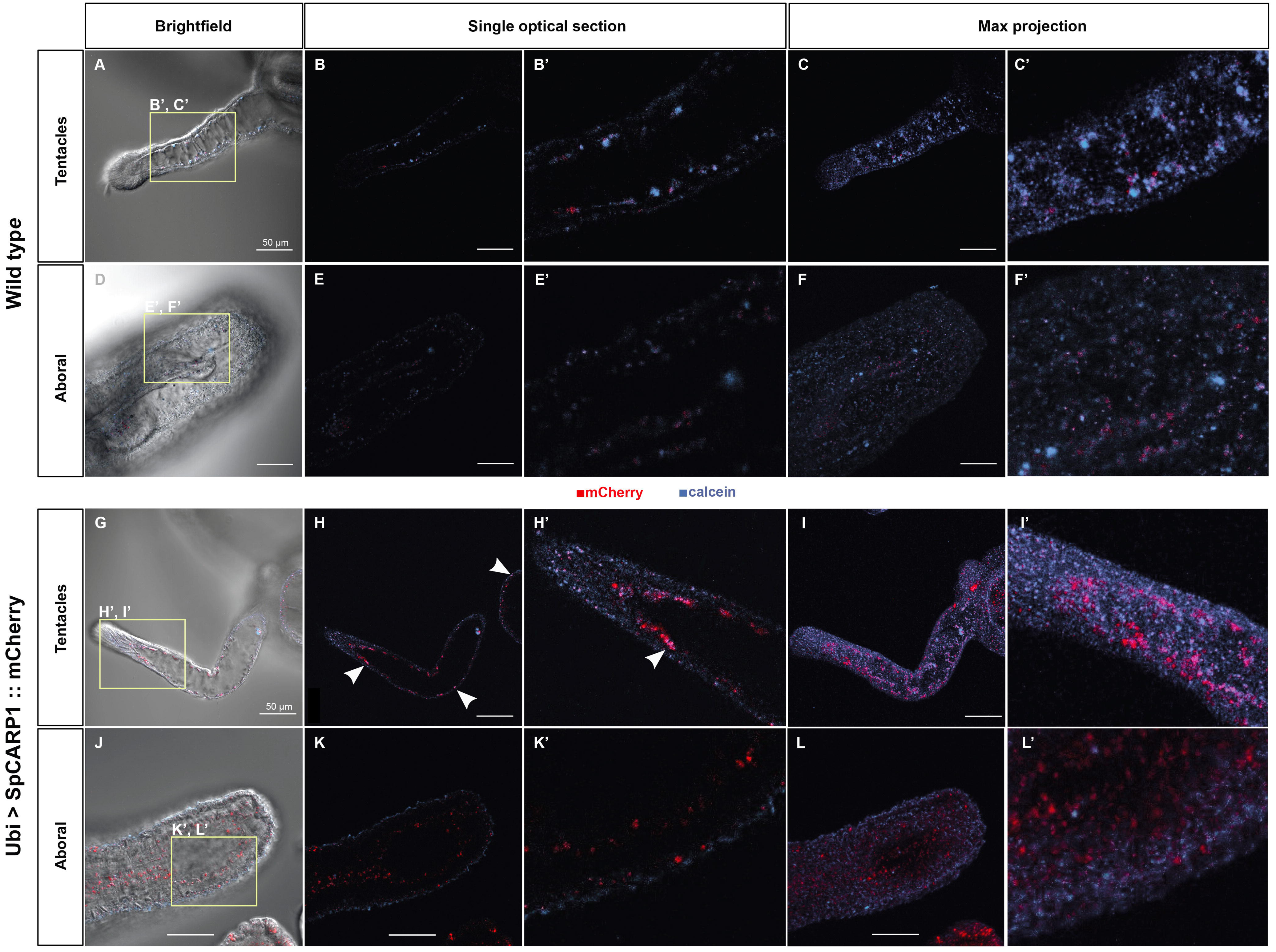
Calcein co-localizes with SpCARP1 in live polyp tentacles (unenriched seawater). Live uninjected polyps show limited calcein signal in tentacular (A–C’) and aboral (D–F’) regions. Live polyps injected with Ubi-CARP1::mCherry plasmid show co-localization of calcein and mCherry signal in tentacular (G–I’) but not aboral (J– L’) regions. Arrowheads indicate co-localization of SpCARP1 with calcein stain. All scale bars = 50μm.

### 3.3 Artificially enriching seawater with carbonate enhances SpCARP1 sequestration of calcium ions

We artificially enriched our seawater with carbonate and/or calcium ions to mimic the concentrated ionic conditions created by coral calcifying cells as they prepare for skeleton deposition. Incubation in carbonate-enriched seawater appears to increase expression in polyp tentacles, as indicated by the expanded expression of mCherry fluorescence in both the endoderm and ectoderm (**Figure 4A–C’**). Such higher expression seems to be accompanied by a significant sequestration of calcium ions (pink reflects the overlap between mCherry and calcein signals in **Figure 4A–C’;** see arrowheads). The body and aboral ends show a similar expression pattern to non-enriched conditions (see **Figure 3J–L’**), with higher mCherry fluorescence in the endoderm and gastrovascular cavity and higher calcein signal in the ectoderm (**Figure 4D–F’**), although some regions of overlapping mCherry-calcein fluorescence are present (see arrowhead in **Figure 4E–E’**). A similar pattern can be observed in polyps with calcium-enriched seawater (**Figure 5A–F’**), although the mCherry signal appears to be dimmer in these conditions compared to carbonate-only-enriched sea water (particularly in the tentacles; see **Figure 5A–C’**). When calcium ions are enriched, mCherry fluorescence is also observable in the ectoderm of the aboral region (**Figures 5D–F’**) compared to just the endoderm of animals in non-enriched solutions (see **Figure 3K–K’**). In addition, compared to the non-enriched conditions (**Figure 3H–I’**), the mCherry signal along the endoderm of the tentacle is less sparse and more homogenous when calcium ions are enriched (**Figure 5B–C’**). Enriching seawater with both calcium and carbonate did not appear to affect the ability of SpCARP1 to sequester and concentrate calcium (**Supplementary Figure 4**).

**Figure 4.**
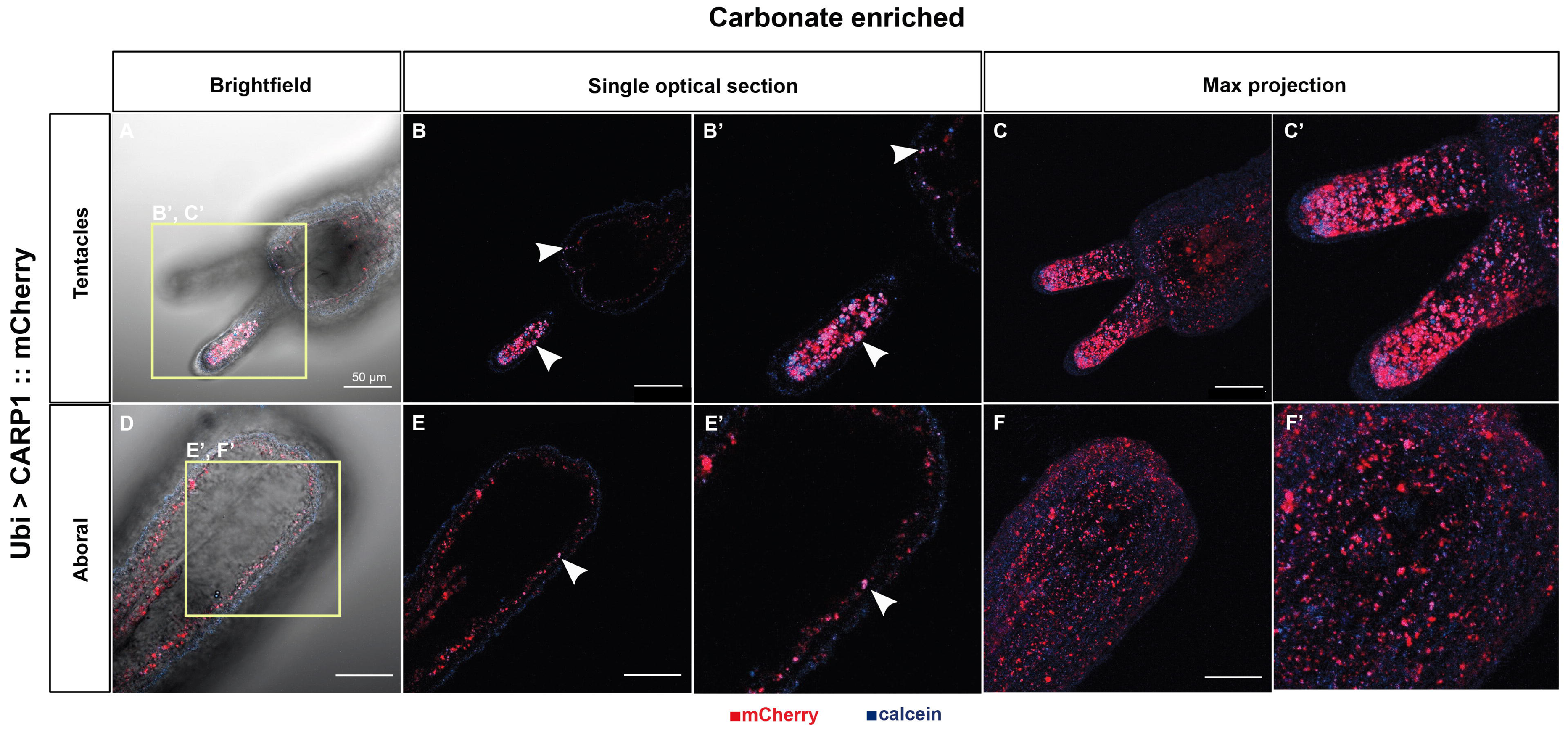
Carbonate-enriched seawater enhances calcium sequestration of SpCARP1 in live polyps. Tentacular (A– C’) and aboral (D–F’) views of live polyps following incubation of carbonate-enriched seawater. White arrowheads indicate co-localization of SpCARP1 with calcein stain. All scale bars = 50μm.

**Figure 5.**
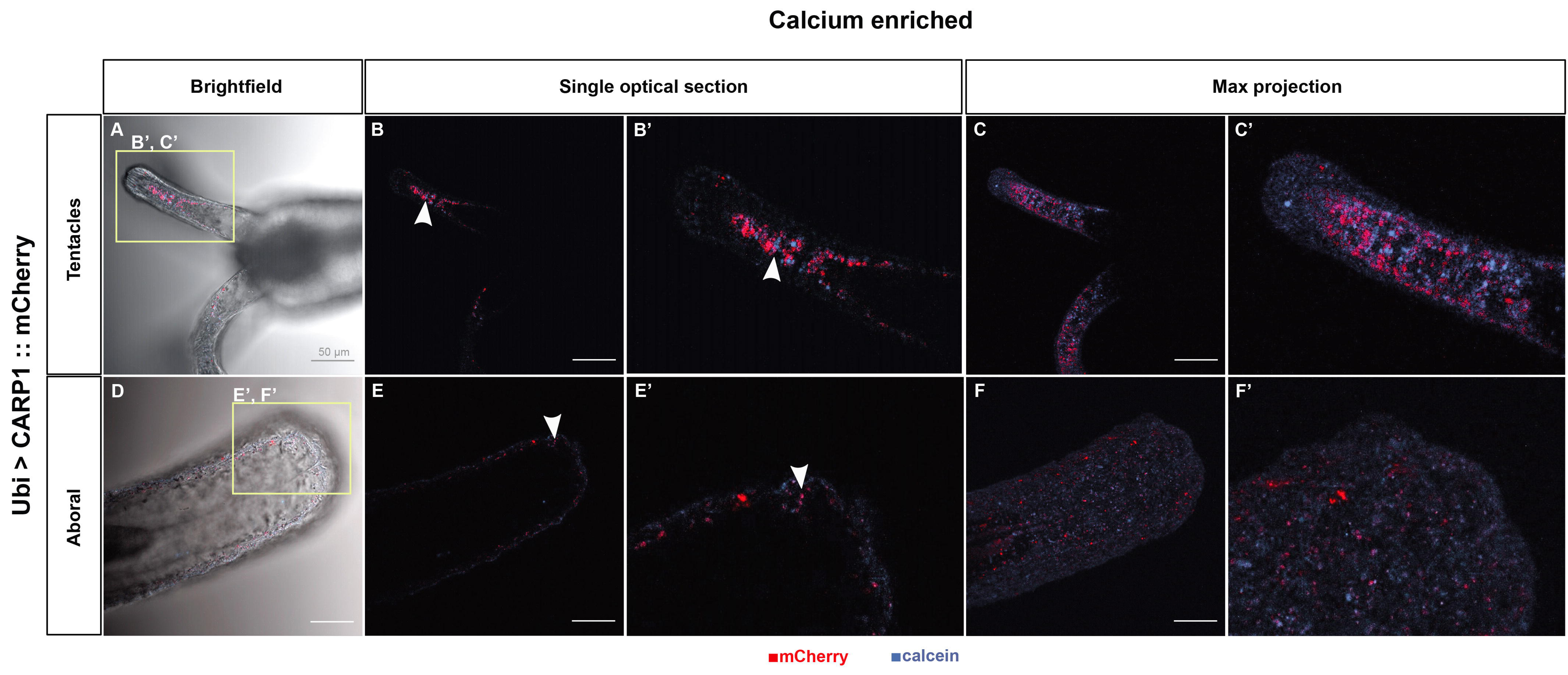
Calcium-enriched seawater does not improve calcium-sequestration of SpCARP1 in live polyps. Tentacular (A–C’) and aboral (D–F’) views of live transgenic polyps following incubation of calcium-enriched seawater. White arrowheads show co-localization of SpCARP1 with calcein stain. All scale bars = 50μm.

## 4 Discussion

We have demonstrated how *N. vectensis*, a soft-bodied anthozoan, may be utilized to study biomineralization *in vivo*.

The putative promoters presented here were selected for optimizing quantity and secretion of target biomineralization IDPs. Ubiquitin, as a regulatory protein that is highly conserved across eukaryotes, should be found in virtually every animal cell. Indeed, the *cis-*regulatory sequence we identified as a ubiquitin promoter appears to drive broad expression of mCherry in a variety of cell types by 24 hpf (**Figure 2A–B**). Such selective expression could be due to an incomplete regulatory sequence or selective protein degradation. Our data shows the putative mucin promoter drives expression of SpCARP1::mCherry within 48 hpf in secretory cells of the aboral ectoderm, with strong mosaic expression throughout the body column and into the tentacles of unfed polyps (**Figure 2C–D**), consistent with the expression of mucin^67^. Mucin-secreting cells are extremely abundant in the aboral ectoderm, and because these animals are excellent regenerators a stable transgenic line with the mucin promoter driving expression of SpCARP1::mCherry should provide abundant material for future analyses of the interactions between SpCARP1 and putative IDPs.

Biomineralizing marine organisms, like corals, have specialized cells that control the chemistry of seawater in a confined space where skeleton deposition occurs, otherwise defined as the “calcifying space.” Corals concentrate calcium and carbonate ions in this calcifying space, and IDPs like SpCARP proteins control the nucleation of aragonite^29^. *N. vectensis*, as a non-calcifying organism, does not possess such specialized calcifying cells. By simulating the biomineralization-favorable conditions of high calcium and high carbonate concentrations, we were able to assess the responsiveness of SpCARP1 and detect regions within *N. vectensis* polyps where biomineralization may be most likely to occur. By supplementing our 1/3X FSW with calcium- and/or carbonate-rich solutions and evaluating calcium sequestration with calcein staining (**Figures 3–5**; see also **Supplementary Figure 4**), we show that the calcium-binding activity of SpCARP1 is primarily concentrated in the tentacles of *N. vectensis* polyps and seems to be enhanced with increased concentrations of carbonate ions in seawater (**Figure 4**), a critical requirement for biomineral nucleation. Our results are consistent with what we would expect from the initial deposition of amorphous calcium carbonate as a precursor to crystalline calcium carbonate. Future studies can further assess the presence of mineral structures in *N. vectensis* tentacles using scanning electron microscopy or polarized light optical microscopy.

We demonstrate that our experimental system is versatile and may be adapted to other forms of biomineralization. We show that *N. vectensis* can express IDPs involved in CaCO_3_ biomineralization of sea urchin spicules and CaPO_4_ precipitation in vertebrate tooth enamel (**Supplementary Figure 2**). The persistence of fluorescent signal in dissociated cells (**Supplementary Figure 3**) suggests it should be possible to isolate and purify proteins for novel uses such as 3D printing of biomineralized material. These results hint at the possibility to expand the use of the *N. vectensis* system to other forms of biomineralization and perhaps even develop novel biomineralized materials for biomedical research.

The primary focus of this study was to show how *N. vectensis* may be utilized to understand the molecular mechanisms that drive coral biomineralization to assist future conservation efforts. This study is the first to attempt to induce biological mineralization in a novel *in vivo* system. We chose the coral acidic protein SpCARP1 because it has been shown to induce rapid mineralization *in vitro*^29^, and to concentrate calcium ions leading to the formation of aragonite crystals in coral proto-polyps derived from cell cultures^68^. However, the calcium ion-concentration activity of such a protein has never been reported in live adult organisms, like we show here in *Nematostella* small polyps.

A single transgenic IDP is likely insufficient to lead to the formation of a mature skeleton. Nevertheless, this study lays the groundwork to establish *N. vectensis* as a tool to interrogate other coral IDPs, transporters, ion pumps, etc. that are implicated in coral biomineralization and that can be coexpressed in the same or adjacent cell types. For example, another coral acid-rich protein, SpCARP4, is of particular interest because it is one of the most abundant proteins in the coral skeleton and has been suggested to guide the formation of calcium carbonate crystals to specific orientations^69^. We predict that *N. vectensis* will be able to tolerate SpCARP4 transgenesis and, if expressed together with SpCARP1, reveal new insights into the interaction between different IDPs and their respective functions in biomineralization. Future studies may help delineate the mechanisms that led calcifying cells to evolve independently in many organisms from a patchwork of nonhomologous proteins and cellular pathways. Such mechanistic studies are necessary to understand how biomineralizing organisms have responded to environmental changes in the past and how they may respond in the future, thereby elucidating how CaCO_3_ biomineralization shapes Earth’s surface environment^47,65,70–72^.

## 5 Conclusion

We demonstrate that *N. vectensis* can both tolerate transgenic expression of intrinsically disordered proteins involved in biomineralization from a range of taxa that can sequester and concentrate calcium ions in a carbonate-enriched seawater solution, providing compelling evidence for the initiation of the biomineralizing process in a non-mineralizing organism. These results highlight the potential of *N. vectensis* in examining the capacity of various cell types to secrete biominerals, opening up opportunities to understand the capacity of cells to acquire novel functions. Our model system may be used as a proxy to coral systems in the lab to test the molecular components of biomineralization that may improve stress tolerance and resilience to native coral populations, thereby filling a much-needed gap in coral research and aiding restoration efforts.

## Competing interests

The authors declare no competing interests.

## Funding

This research was funded by NSF grant # IOS 2314456.

## Notes

### Competing Interest Statement

The authors have declared no competing interest.

## References

1. Reaka-Kudla, M. L. Biodiversity II: Understanding and Protecting Our Biological Resources. (Joseph Henry Press, 1997).

2. Small, A. M., Adey, W. H. & Spoon, D. Are Current Estimates of Coral Reef Biodiversity Too Low? the View through the Window of a Microcosm. Atoll Res. Bull. 1–20 (1998) doi:10.5479/SI.00775630.458.1.

3. Wagner, D. et al. Coral Reefs of the High Seas: Hidden Biodiversity Hotspots in Need of Protection. Front. Mar. Sci. 7, 567428 (2020).

4. Arkema, K. K. et al. Coastal habitats shield people and property from sea-level rise and storms. Nat. Clim. Change 3, 913–918 (2013).

5. Moberg, F. & Folke, C. Ecological goods and services of coral reef ecosystems. Ecol. Econ. 29, 215–233 (1999).

6. Baker, A. C., Glynn, P. W. & Riegl, B. Climate change and coral reef bleaching: An ecological assessment of long-term impacts, recovery trends and future outlook. Estuar. Coast. Shelf Sci. 80, 435–471 (2008).

7. Addadi, L., Joester, D., Nudelman, F. & Weiner, S. Mollusk Shell Formation: A Source of New Concepts for Understanding Biomineralization Processes. Chem Eur J 12, 980–987 (2006).

8. Gotliv, B. A., Addadi, L. & Weiner, S. Mollusk Shell Acidic Proteins: In Search of Individual Functions. ChemBioChem 4, 522–529 (2003).

9. Marie, B. et al. Different secretory repertoires control the biomineralization processes of prism and nacre deposition of the pearl oyster shell. (2012) doi:10.1073/pnas.1210552109.

10. Weiner, S. Biomineralization: A structural perspective. J. Struct. Biol. 163, 229–234 (2008).

11. Decker, G. L., Morrill, J. B. & Lennarz, W. J. Characterization of sea urchin primary mesenchyme cells and spicules during biomineralization in vitro. Development 101, 297–312 (1987).

12. Gildor, T., Winter, M. R., Layous, M., Hijaze, E. & Ben-Tabou de-Leon, S. The biological regulation of sea urchin larval skeletogenesis – From genes to biomineralized tissue. J. Struct. Biol. 213, (2021).

13. Politi, Y., Arad, T., Klein, E., Weiner, S. & Addadi, L. Sea Urchin Spine Calcite Forms via a Transient Amorphous Calcium Carbonate Phase. New Ser. 306, 1161–1164 (2004).

14. Mor Khalifa, G., Levy, S. & Mass, T. The calcifying interface in a stony coral primary polyp: An interplay between seawater and an extracellular calcifying space. J. Struct. Biol. 213, 107803 (2021).

15. Neder, M. et al. Mineral formation in the primary polyps of pocilloporoid corals. Acta Biomater. 96, 631–645 (2019).

16. Von Euw, S. et al. Biological control of aragonite formation in stony corals. Science 356, 933–938 (2017).

17. Kahil, K. et al. Cellular pathways of calcium transport and concentration toward mineral formation in sea urchin larvae. Proc. Natl. Acad. Sci. U. S. A. 117, 30957–30965 (2020).

18. Kahil, K., Weiner, S., Addadi, L. & Gal, A. Ion Pathways in Biomineralization: Perspectives on Uptake, Transport, and Deposition of Calcium, Carbonate, and Phosphate. Cite This J Am Chem Soc 143, (2021).

19. Bose, H. & Satyanarayana, T. Microbial Carbonic Anhydrases in Biomimetic Carbon Sequestration for Mitigating Global Warming: Prospects and Perspectives. Front. Microbiol. 0, 1615 (2017).

20. Le Roy, N., Jackson, D. J., Marie, B., Ramos-Silva, P. & Marin, F. The evolution of metazoan a_α_-carbonic anhydrases and their roles in calcium carbonate biomineralization. Front. Zool. 11, 1–16 (2014).

21. Moya, A. et al. Carbonic Anhydrase in the Scleractinian Coral Stylophora pistillata: CHARACTERIZATION, LOCALIZATION, AND ROLE IN BIOMINERALIZATION *. J. Biol. Chem. 283, 25475–25484 (2008).

22. Voigt, O., Adamski, M., Sluzek, K. & Adamska, M. Calcareous sponge genomes reveal complex evolution of _α_-carbonic anhydrases and two key biomineralization enzymes. BMC Evol. Biol. 14, 1–19 (2014).

23. Addadi, L., Raz, S. & Weiner, S. Taking advantage of disorder: Amorphous calcium carbonate and its roles in biomineralization. Adv. Mater. 15, 959–970 (2003).

24. Nebel, H., Neumann, M., Mayer, C. & Epple, M. On the Structure of Amorphous Calcium Carbonate—A Detailed Study by Solid-State NMR Spectroscopy. Inorg. Chem. 47, 7874–7879 (2008).

25. Whiticar, M. J., Suess, E., Wefer, G. & Müller, P. J. Calcium Carbonate Hexahydrate (Ikaite): History of Mineral Formation as Recorded by Stable Isotopes. Minerals 12, 1627 (2022).

26. Mass, T. et al. Amorphous calcium carbonate particles form coral skeletons. Proc. Natl. Acad. Sci. U. S. A. 114, E7670–E7678 (2017).

27. Gower, L. B. & Odom, D. J. Deposition of calcium carbonate films by a polymer-induced liquid-precursor (PILP) process. J. Cryst. Growth 210, 719–734 (2000).

28. George, A., Sabsay, B., Simonian, P. A. L. & Veiss, A. THE JOURNAL OF BIOLOGICAL CHEMISTRY Characterization of a Novel Dentin Matrix Acidic Phosphoprotein IMPLICATIONS FOR INDUCTION OF BIOMINERALIZATION*. J. Biol. Chem. 268, 12624–12630 (1993).

29. Mass, T. et al. Cloning and characterization of four novel coral acid-rich proteins that precipitate carbonates in vitro. Curr. Biol. 23, 1126–1131 (2013).

30. Helman, Y. et al. Extracellular matrix production and calcium carbonate precipitation by coral cells in vitro. Proc. Natl. Acad. Sci. 105, 54–58 (2008).

31. Veis, A. & Dorvee, J. R. Biomineralization mechanisms: A new paradigm for crystal nucleation in organic matrices. Calcif. Tissue Int. 93, 307–315 (2013).

32. Wilt, F. H., Killian, C. E., Hamilton, P. & Croker, L. The dynamics of secretion during sea urchin embryonic skeleton formation. Exp. Cell Res. 314, 1744–1752 (2008).

33. Kalmar, L., Homola, D., Varga, G. & Tompa, P. Structural disorder in proteins brings order to crystal growth in biomineralization. Bone 51, 528–534 (2012).

34. Moradian-Oldak, J. & George, A. Biomineralization of Enamel and Dentin Mediated by Matrix Proteins. J. Dent. Res. 100, 1020–1029 (2021).

35. Ndao, M. et al. Intrinsically disordered mollusk shell prismatic protein that modulates calcium carbonate crystal growth. Biomacromolecules 11, 2539–2544 (2010).

36. Rose-Martel, M., Smiley, S. & Hincke, M. T. Novel identification of matrix proteins involved in calcitic biomineralization. J. Proteomics 116, 81–96 (2015).

37. Gorski, J. P. Calcified Tissue International Acidic Phosphoproteins from Bone Matrix: A Structural Rationalization of Their Role in Biomineralization. Calcif Tissue Int 50, 391–396 (1992).

38. Gotliv, B. A. et al. Asprich: A novel aspartic acid-rich protein family from the prismatic shell matrix of the bivalve Atrina rigida. Chembiochem Eur. J. Chem. Biol. 6, 304–314 (2005).

39. Moradian-Oldak, J., Frolow, F., Addadi, L. & Weiner, S. Interactions between acidic matrix macromolecules and calcium phosphate ester crystals: relevance to carbonate apatite formation in biomineralization. Proc. R. Soc. Lond. B Biol. Sci. 247, 47–55 (1992).

40. Suzuki, M. et al. An Acidic Matrix Protein, Pif, Is a Key Macromolecule for Nacre Formation. (2009) doi:10.1126/science.1173793.

41. Kabakoff, B., Hwang, S. P. L. & Lennarz, W. J. Characterization of post-translational modifications common to three primary mesenchyme cell-specific glycoproteins involved in sea urchin embryonic skeleton formation. Dev. Biol. 150, 294–305 (1992).

42. Dhami, N. K., Reddy, M. S. & Mukherjee, A. Synergistic role of bacterial urease and carbonic anhydrase in carbonate mineralization. Appl. Biochem. Biotechnol. 172, 2552–2561 (2014).

43. Li, W. et al. Calcium carbonate precipitation and crystal morphology induced by microbial carbonic anhydrase and other biological factors. Process Biochem. 45, 1017–1021 (2010).

44. Smith, K. S. & Ferry, J. G. Prokaryotic carbonic anhydrases. FEMS Microbiol. Rev. 24, 335–366 (2000).

45. Ettensohn, C. A. & Malinda, K. M. Size regulation and morphogenesis: A cellular analysis of skeletogenesis in the sea urchin embryo. Development 119, 155–167 (1993).

46. Beniash, E., Addadi, L. & Weiner, S. Cellular Control Over Spicule Formation in Sea Urchin Embryos: A Structural Approach. J. Struct. Biol. 125, 50–62 (1999).

47. Clark, M. S. Molecular mechanisms of biomineralization in marine invertebrates. J. Exp. Biol. 223, (2020).

48. Wang, X. et al. The Evolution of Calcification in Reef-Building Corals. Mol. Biol. Evol. 38, 3543–3555 (2021).

49. Renfer, E. & Technau, U. Meganuclease-assisted generation of stable transgenics in the sea anemone Nematostella vectensis. Nat. Protoc. 12, 1844–1854 (2017).

50. Layden, M. J., Röttinger, E., Wolenski, F. S., Gilmore, T. D. & Martindale, M. Q. Microinjection of mRNA or morpholinos for reverse genetic analysis in the starlet sea anemone, Nematostella vectensis. Nat. Protoc. 8, 924–934 (2013).

51. Hand, C. & Uhlinger, K. R. The Culture, Sexual and Asexual Reproduction, and Growth of the Sea Anemone Nematostella vectensis. Biol. Bull. 182, 169–176 (1992).

52. Fritzenwanker, J. H. & Technau, U. Induction of gametogenesis in the basal cnidarian Nematostella vectensis(Anthozoa). Dev. Genes Evol. 212, 99–103 (2002).

53. Chin, J. X., Chung, B. K. S. & Lee, D. Y. Codon Optimization OnLine (COOL): A web-based multi-objective optimization platform for synthetic gene design. Bioinformatics 30, 2210–2212 (2014).

54. Koressaar, T. et al. Primer3_masker: integrating masking of template sequence with primer design software. Bioinforma. Oxf. Engl. 34, 1937–1938 (2018).

55. Hernández, G., Osnaya, V. G. & Pérez-Martínez, X. Conservation and Variability of the AUG Initiation Codon Context in Eukaryotes. Trends Biochem. Sci. 44, 1009–1021 (2019).

56. Waldo, G. S., Standish, B. M., Berendzen, J. & Terwilliger, T. C. Rapid protein-folding assay using green fluorescent protein. Nat. Biotechnol. 17, 691–695 (1999).

57. Almagro Armenteros, J. J. et al. SignalP 5.0 improves signal peptide predictions using deep neural networks. Nat. Biotechnol. 2019 374 37, 420–423 (2019).

58. Putnam, N. H. et al. Sea anemone genome reveals ancestral eumetazoan gene repertoire and genomic organization. Science 317, 86–94 (2007).

59. Marlow, H. Q., Srivastava, M., Matus, D. Q., Rokhsar, D. & Martindale, M. Q. Anatomy and development of the nervous system of Nematostella vectensis, an anthozoan cnidarian. Dev. Neurobiol. 69, 235–254 (2009).

60. Marlow, H., Roettinger, E., Boekhout, M. & Martindale, M. Q. Functional roles of Notch signaling in the cnidarian Nematostella vectensis. Dev. Biol. 362, 295–308 (2012).

61. Schindelin, J. et al. Fiji: an open-source platform for biological-image analysis. Nat. Methods 2012 97 9, 676–682 (2012).

62. Pierrot, D., Lewis, E. & Wallace, D. MS Excel Program Developed for CO2 System Calculations. https://marine.gov.scot/sma/content/ms-excel-program-developed-co2-system-calculations (2006).

63. Mehrbach, C., Culberson, C. H., Hawley, J. E. & Pytkowicx, R. M. MEASUREMENT OF THE APPARENT DISSOCIATION CONSTANTS OF CARBONIC ACID IN SEAWATER AT ATMOSPHERIC PRESSURE1. Limnol. Oceanogr. 18, 897–907 (1973).

64. Dickson, A. G. & Millero, F. J. A comparison of the equilibrium constants for the dissociation of carbonic acid in seawater media. Deep Sea Res. Part Oceanogr. Res. Pap. 34, 1733–1743 (1987).

65. Gilbert, P. U. P. A. et al. Biomineralization: Integrating mechanism and evolutionary history. Sci. Adv. 8, (2022).

66. Sebé -Pedró, A., Saudemont, B., Spitz, O., Tanay, A. & Marlow, H. Cnidarian Cell Type Diversity and Regulation Revealed by Whole-Organism Single-Cell RNA-Seq In Brief. (2018) doi:10.1016/j.cell.2018.05.019.

67. Steinmetz, P. R. H., Aman, A., Kraus, J. E. M. & Technau, U. Gut-like ectodermal tissue in a sea anemone challenges germ layer homology. Nat. Ecol. Evol. 1, 1535–1542 (2017).

68. Mass, T., Drake, J. L., Heddleston, J. M. & Falkowski, P. G. Nanoscale Visualization of Biomineral Formation in Coral Proto-Polyps. (2017) doi:10.1016/j.cub.2017.09.012.

69. Mass, T., Drake, J. L., Peters, E. C., Jiang, W. & Falkowski, P. G. Immunolocalization of skeletal matrix proteins in tissue and mineral of the coral Stylophora pistillata. Proc. Natl. Acad. Sci. U. S. A. 111, 12728–12733 (2014).

70. Drake, J. L. et al. How corals made rocks through the ages. Glob. Change Biol. 26, 31–53 (2020).

71. Hönisch, B. et al. The geological record of ocean acidification. Science 335, 1058–1063 (2012).

72. Kawahata, H. et al. Perspective on the response of marine calcifiers to global warming and ocean acidification—Behavior of corals and foraminifera in a high CO2 world “hot house”. Prog. Earth Planet. Sci. 2019 61 6, 1–37 (2019).

