## Supplementary Figures for "A novel *in vivo* system to study coral biomineralization in the starlet sea anemone (*Nematostella vectensis*)"

| **A**  **pCS2+8CmCherry** (Addgene Plasmid #34935)  4848 bp  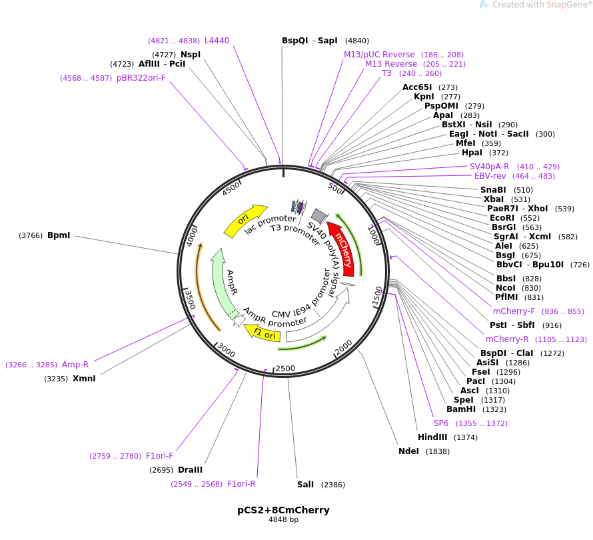 | **B**  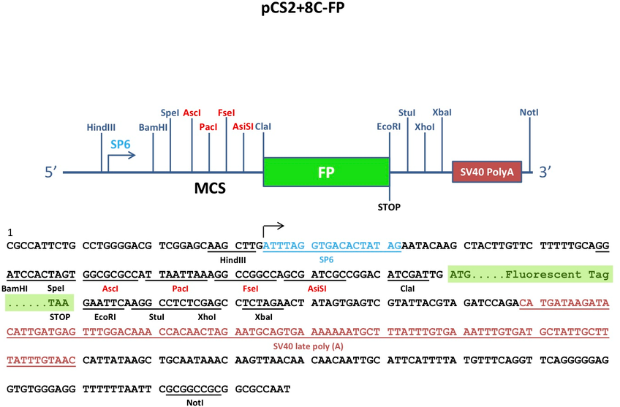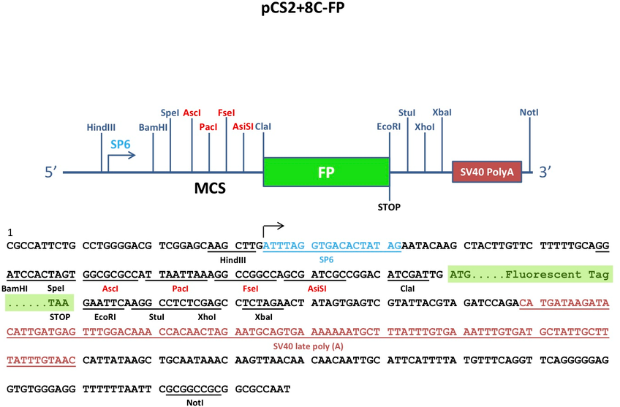  ORF  Transgene cassette  Promoter |
| --- | --- |
| 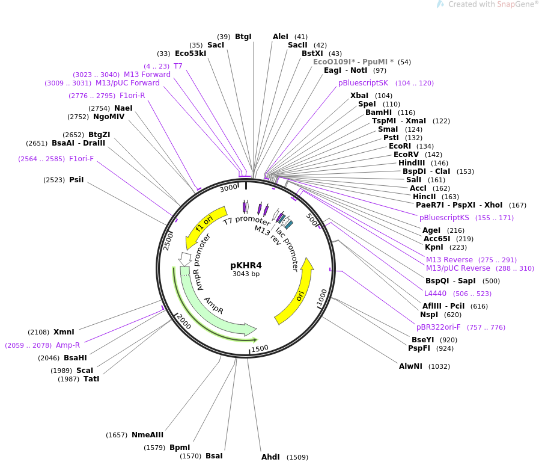  **C**  **pKHR4** (Addgene #74592)  3043 bp | 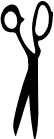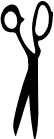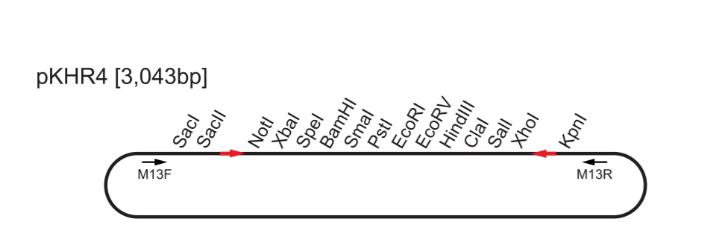  I-SceI  Transgene cassette insertion site  **D**  185 . . . . . . . . . . . . . . . 71  I-SceI |

**Supplementary Figure 1.** *Transferring the promoter-ORF mCherry fragment from PCS2 into the pKHR4 vector.* Vector map of pCS2+8CmCherry (A) and transgene cassette incorporating *Nv* promoters, Open Reading Frames (ORFs), fluorescent protein, and SV40 PolyA tail (B). Promoters were cloned in frame with SpeI and AscI restriction sites. ORFs were cloned in frame with AscI and ClaI restriction sites. The transgene cassette was cut with SpeI and NotI restriction enzymes and cloned into the pKHR4 vector (C–D). MCS = Multiple Cloning Site. FP = Fluorescent Protein.

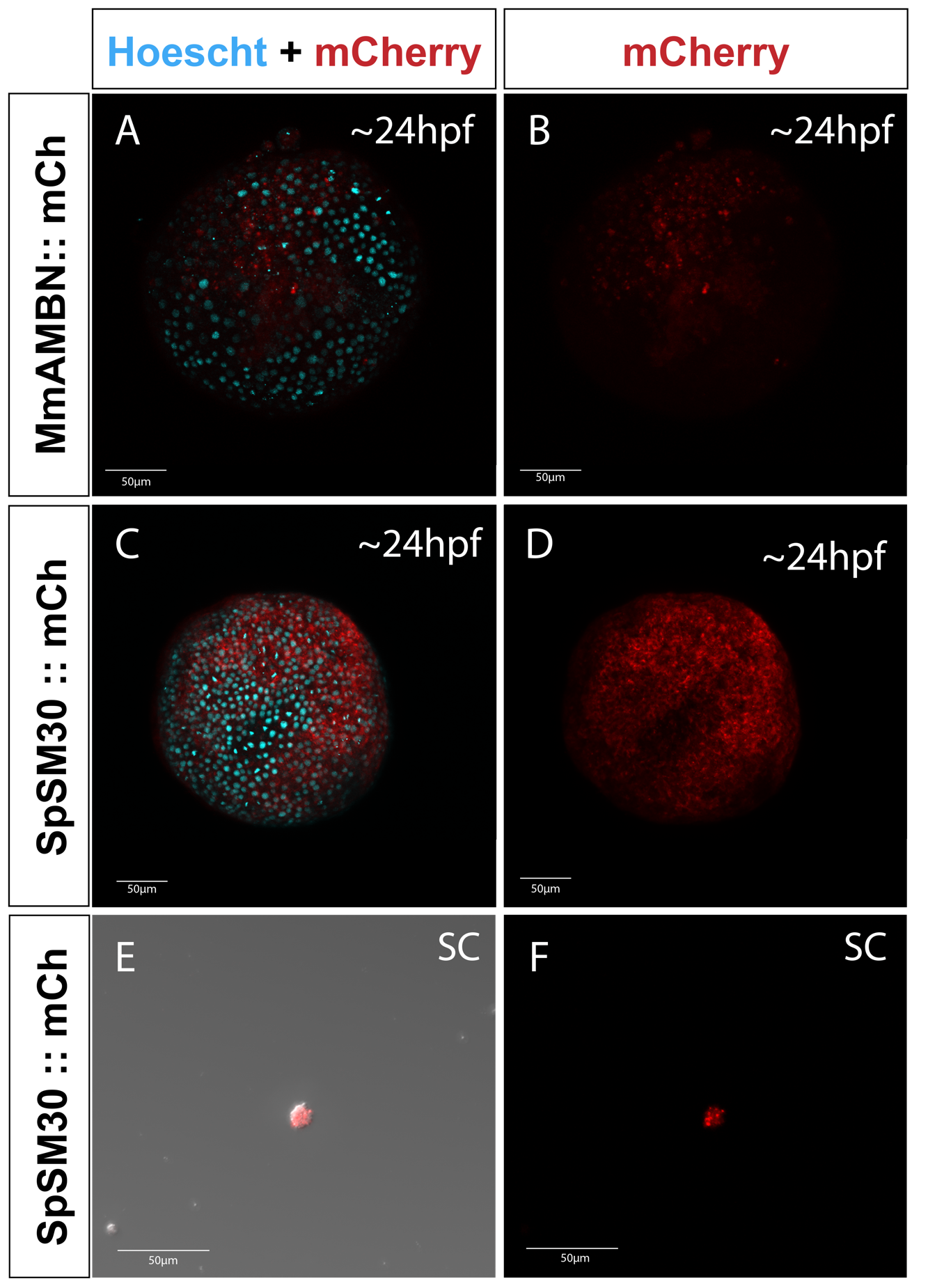
**Supplementary Figure 2.** *Injection of mRNA biomineralization IDPs are toxic*. Animals microinjected with mRNA for AMBN::mCh (A–B) and SM30::mCh (C–D) show broad expression within 24 hpf. Dissociated single cells also express fluorescent signal (E–F).

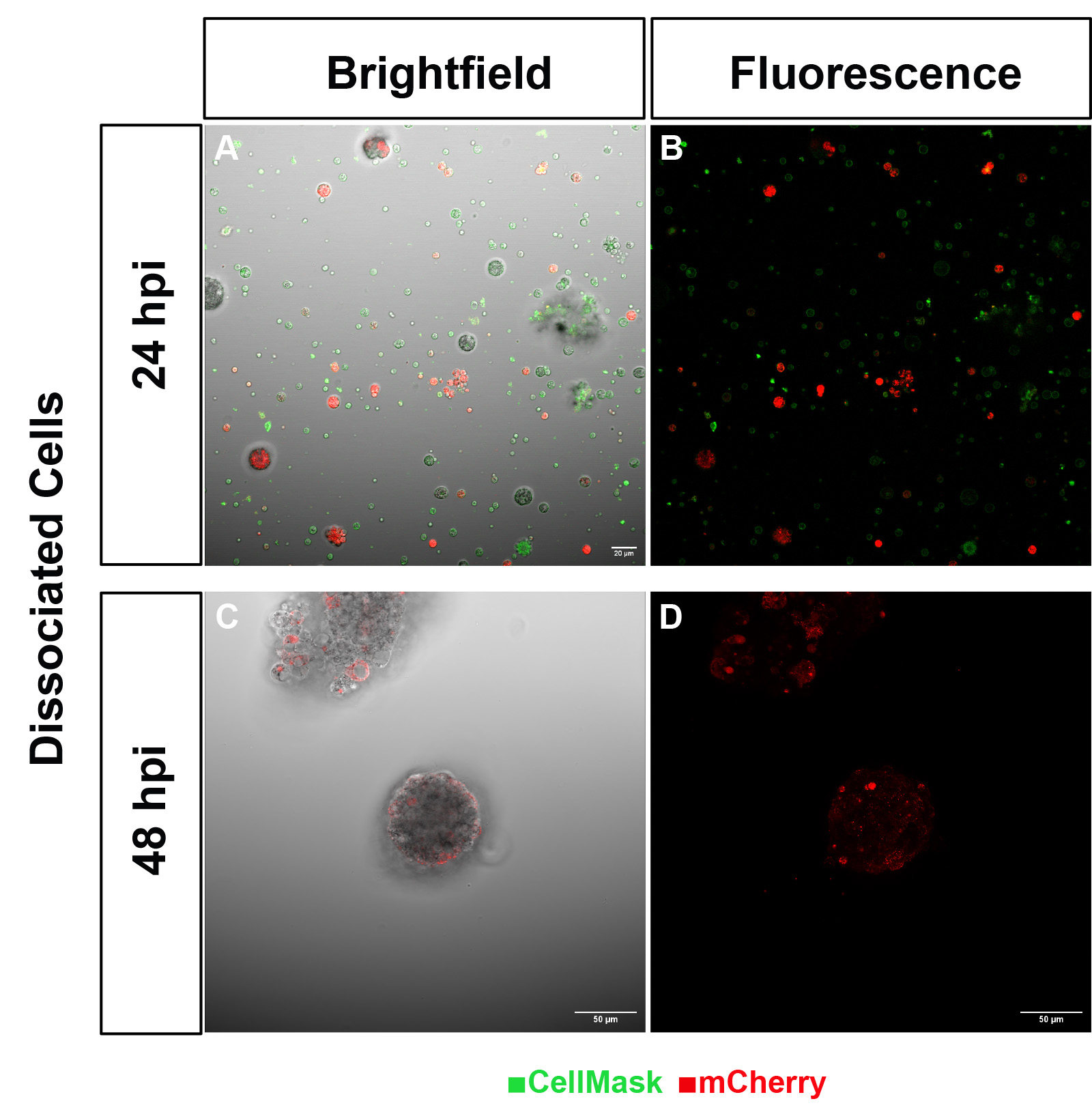

**Supplementary Figure 3.** *CARP1::mCherry signal persists in dissociated cells 48 hours post injection (hpi)*. Dissociated cells express CARP1::mCherry (A–D) and may be used for purification or further culturing. Cells form aggregates 24 h after dissociation (C–D).

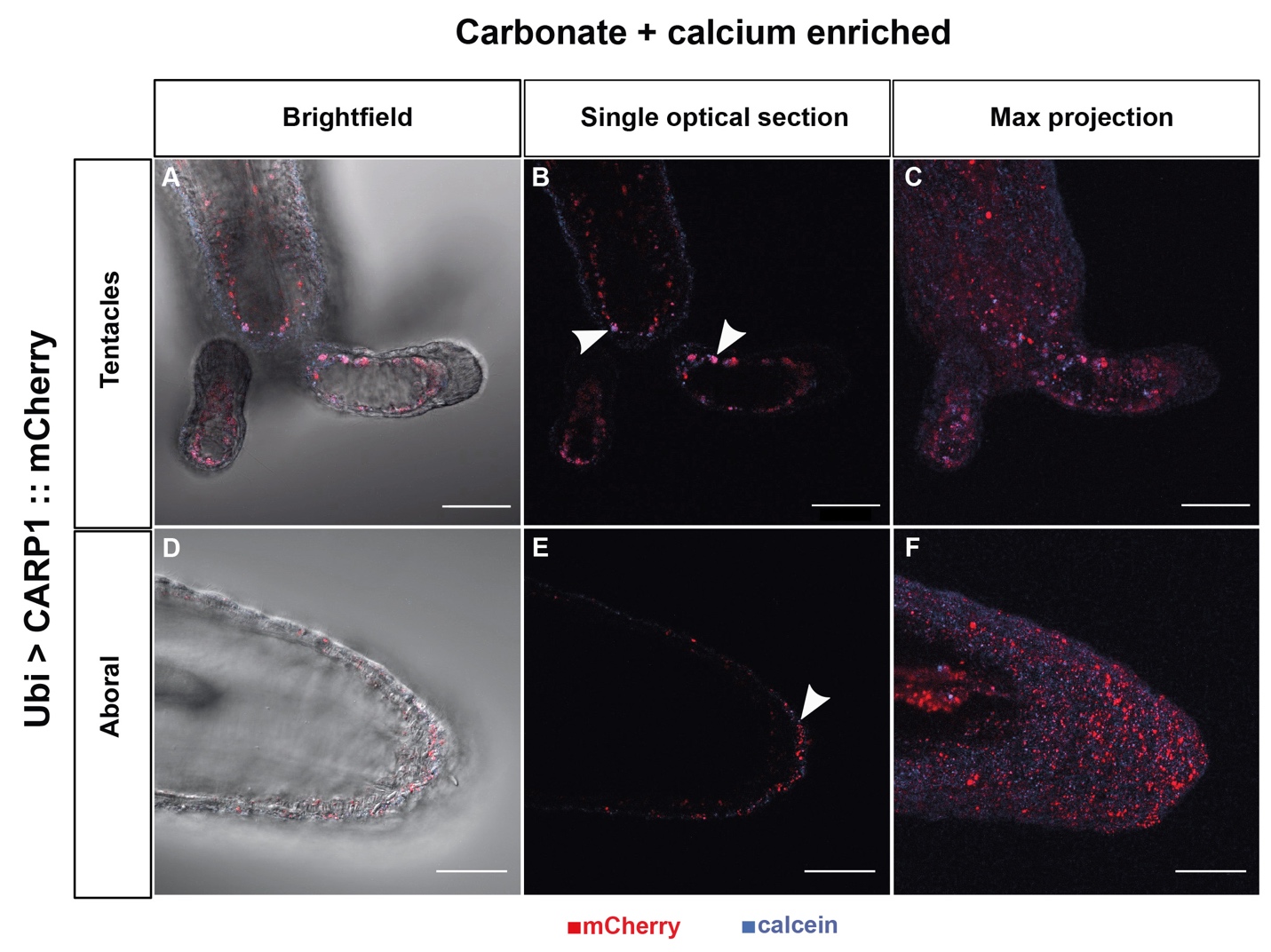

**Supplementary Figure 4.** *Calcium- and Carbonate-enriched seawater does not improve calcium sequestration of CARP1 in live polyps.* Oral (A–C) and aboral (D–F) views. White arrows indicate co-localization of CARP1 and calcein stain. All scalebars = 50µm.
