## Supplementary Table 1 for "A novel *in vivo* system to study coral biomineralization in the starlet sea anemone (*Nematostella vectensis*)"

| Gene | Protein ID | Size (kb) | Nucleotide Seq | F Primer | R Primer |
| --- | --- | --- | --- | --- | --- |
| SpCARP1 | KC148537 | 1.041 | ATGTTTCATTCTTGGTGGATGACGCTATTGATTTTAGGTAGCACCGTGAGTTTTGTTTTCACGGAAGGTGATCATTTAAAGCCTGGACACTCAGAAGATGAACACGACGAAGACGAACATGATGAAGAAATGGCAGACCATGCAGACGAACAGAATCCTGCCGATGAGGAAGAAACAGAGGATGAGGAAAAAGACGACGATAAAATGGAAGATGACAGCGATGATGACGAAGAAGATGAAAGTCAAGGAGATGATGAAGGAGAAGATGAGAATGATCAAAGTCACCTCGAACACGACGCATTTTTAGGAGACAACTACACTGAATTCAAAGGGCTTTCCCCCGAGGACGCAAAAACCAAACTGGCACAACTAATCAAAGATGAGGTGGATTTGAACAAGGACGGAGTGTTGACAGAGGATGAGATCAGACAGCGATTCCACGTAACTACAAAAGAATACAGGAAAAAAGAAGTCATGGAAACGATGAAGCAACATGATGAGGACAAAGATGGGAAAGTATCGTGGGAAGAATTTAAAAAAGGTCATTTCTCTGATGATGGCAAAGATGAAGATGCTAAAGAACAAATGAAAGAAGATGAAGAAAAATTCAAATTTGCTGATGAAGATGGAGATGGAAAGTTAGACTTGGAGGAATATATGGCGTTTTACCACCCAGGGGATAATCCAAGAATGACGGAGTTCACCATAGAAGATAGCCTCAAAAAGCATGACAAGGATAAGGATGGTCAAGTATCAAAGAAAGAATTTTTAGCTACATTCAGCGATGTAAATGATGACGCCAAAGAAGAAATGGAAAAGGATTTTAATAACAACTTTGATAAGGACAAGAATGGTAGATTAAACAAAGAAGAAATGAAAAGCTGGCTGTTCCCAGATGACGACTTCTCCACTGAAGAACCTAAAACACTCATCAAAGAAGCTGATGAAGACAAAGATGGCAAACTCACCATGGATGAGATAATGAAAAATTATAAAGTCTTCATCGAGGATGAACCTGAAGATTCAAGCCACGATGAATTA | NA | NA |
| SpSM30 | NP_999766.1 | 0.87 | ATGAGGGGATTTGTTTATGTGCTGGTGTGTGTTCTTGCTCTTGCATCATTTTCAAGAGCACAGCTTCCTGGTGCAGGAGGACCAGTTTTGCCAGGTGGTGGACCCACCATTGGTCCTGTGAATCCAGACCCAACAAGAACAGAAGTTTGTGCAAAGTTCTGGGTGCAAGAAGGAAATTCTTGTTATTTGTTTGACAGCGGAGCATTTCTTAGACAAGTTGCCGCCTCAAGGCCAGTGGTGGTCAATAATCAAGATGGTCTGTTCCAGGCTGCTGCAAACATGTACTGTGGACAAATGCACCCAAATGCAAGCCTGGTCACTGTCAACAGCCTTGAAGAAAACAATTTTTTGTATGAATGGGCAGTGCGCATGATGATCGAGCCAGAGCCTGTTTGGATTGGCCTTCATGCTGGACCTACTGGACAGTGGCAATGGTACAGTGGAGAACCTGTCACATACACAAACTGGGAAAGGATGACCCCTCCTATTGCAGAACCAGGACTTGGTGCCATGATTTTTGATGCTGACATCATCGCTCAGATGTTCAACAACCAAGTGGAGATCACACCACAGTGGGTTCCAGAACAAGCCATCAATGACAGACATGCACTTATTTGTGAGTACCATCCATCTGGAATGACAGCAGCCGCTGCCGCACCTACAAATGCTCCCACATTTCCACCAATGACTACAGCTCCACCCATGGCAGCAACAACAAGGGGTCCAGTCATGTTTCAAAATAACCCAAGAAATCTTGTGAACAGTTTGACAGGAGGGAGATTTGGAGGCTCTCTTCTTCACGAGATTCCTAGAAGACAGAGAATGAGACCAAGCAACTACAGAAAAAACCCTTATTTTGGTATCCAGCCA | CAGGCGCGCCATGAGGGGATTTGTTTATGTG | CCATCGATTGGCTGGATACCAAAATAAGGGTTT |
| MmAMBN | NM_001303431.1 | 1.269 | ATGTCAGCATCTAAGATTCCACTTTTCAAAATGAAGGGCCTGATCCTGTTCCTGTCCCTAGTGAAAATGAGCCTCGCCGTGCCGGCATTTCCTCAACAACCTGGGGCTCAAGGCATGGCACCTCCTGGCATGGCTAGTTTGAGCCTTGAGACAATGAGACAGTTGGGAAGCTTGCAGGGACTCAACGCACTTTCTCAGTATTCTAGACTTGGCTTTGGAAAAGCACTTAATAGTTTATGGTTGCACGGACTTCTCCCACCGCATAACTCTTTCCCATGGATAGGACCAAGGGAACATGAAACCCAGCAGTATGAATATTCTTTGCCTGTGCATCCCCCACCTCTCCCATCACAGCCATCCTTGCAGCCTCACCAGCCAGGACTGAAACCCTTCCTCCAGCCCACTGCTGCAACCGGTGTCCAGGTCACACCCCAGAAGCCAGGGCCTCAGCCTCCAATGCACCCTGGACAGCTGCCCTTGCAGGAAGGAGAGCTGATAGCACCAGATGAGCCGCAGGTGGCACCATCCGAAAACCCACCAACACCTGAGGTACCAATAATGGATTTTGCTGATCCACAATTTCCAACCGTGTTCCAGATCGCCCGTTCAATATCTCGGGGACCAATGGCACACAACAAAGCATCCGCTTTTTACCCAGGAATGTTTTACATGTCTTATGGAGCAAACCAATTGAATGCTCCTGCCAGAATTGGCTTCATGAGTTCAGAAGAAATGCCTGGAGAAAGAGGAAGTCCCATGGCCTATGGAACTCTGTTCCCAAGATTTGGAGGCTTCAGGCAAACCCTTAGGAGACTGAATCAGAATTCACCCAAGGGAGGAGACTTTACTGTGGAAGTAGATTCCCCAGTATCTGTTACCAAAGGCCCTGAAAAAGGAGAAGGTCCAGAAGGCTCTCCACTGCAAGAGGCCAACCCAGGCAAACGGGAAAACCCCGCTCTCCTTTCACAAATGGCACCTGGGGCCCATGCAGGACTTCTTGCTTTCCCCAATGACCACATCCCCAGTATGGCAAGGGGTCCTGCAGGGCAAAGACTCCTTGGAGTCACCCCTGCAGCTGCAGACCCACTGATCACCCCTGAATTAGCAGAAGTTTATGAAACCTATGGTGCTGATGTTACCACACCCCTGGGTGATGGAGAAGCAACCATGGATATCACCATGTCCCCAGACACTCAGCAGCCACTGCTACCTGGAAACAAAGTGCACCAGCCCCAGGTGCACAACGCATGGCGTTTCCAAGAGCCCTGA | AAGGCGCGCCAAATTAATACGA | CCATCGATCGGGCTCTTGGAAA |
