## Supplementary Table 2 for "A novel *in vivo* system to study coral biomineralization in the starlet sea anemone (*Nematostella vectensis*)"

**Supplementary Figure 2**

| Gene Promoter | ID | Scaffold | Start Coordinate | End Coordinate | Size (kb) | F Primer with PacI site | R Primer with AscI site |
| --- | --- | --- | --- | --- | --- | --- | --- |
| Mucin1.8 | v1g203270 | 41 | 1082156 | 1080338 | 1.8 | TTAATTAAGCCCCCGGGCCATTTTGGG | GGCGCGCCCTTACGGAAAAAATAT |
| Mucin2.8 | v1g203270 | 41 | 1083139 | 1080338 | 2.8 | TTAATTGGCAGCTATTCCCCCACTTGC | GGCGCGCCCTTACGGAAAAAATAT |
| Ubi3.5 | v1g217964 | 320 | 94027 | 97431 | 3.5 | CCTTAATTAAAGGGTTTACGGGTCCTCTTG | GAGGCGCGCCTGAAAAGGCGGAATATCGGG |
