## Supplementary Table 3 for "A novel *in vivo* system to study coral biomineralization in the starlet sea anemone (*Nematostella vectensis*)"

| pH | pCO_2_ (µatm) | HCO_3_^-^ (µmol/kg·SW) | CO_3_^2-^ (µmol/kg·SW) | ΩCa | ΩAr |
| --- | --- | --- | --- | --- | --- |
| 7.900 ± 0.006 | 500.700 ± 0.006 | 845.900 ± 1.174 | 21.800 ± 0.387 | 0.639 ± 0.010 | 0.371 ± 0.006 |
| 8.100 ± 0.003 | 2695.5 ± 35.012 | 7342.700 ± 10.374 | 318.900 ± 4.839 | 9.146 ± 0.109 | 5.386 ± 0.076 |
| 7.800 ± 0.006 | 585.200 ± 11.470 | 849.900 ± 1.141 | 19.600 ± 0.462 | 0.563 ± 0.012 | 0.331 ± 0.008 |
| 8.100 ± 0.003 | 2719.500 ± 6.393 | 7665.600 ± 3.601 | 355.800 ± 1.542 | 10.037 ± 0.016 | 5.978 ± 0.022 |
